# The impact of bilingualism on executive functions and working memory in young adults

**DOI:** 10.1101/449827

**Authors:** Eneko Antón, Manuel Carreiras, Jon Andoni Duñabeitia

**Affiliations:** Facultad de Lenguas y Educación, Universidad Nebrija; Madrid, Spain; BCBL. Basque Center on Cognition, Brain and Language; Donostia, Spain.; Ikerbasque, Basque Foundation for Science; Bilbao, Spain.; Euskal Herriko Unibertsitatea – Universidad del País Vasco; Bilbao, Spain.

**Keywords:** Bilingualism, executive functioning, working memory

## Abstract

A bilingual advantage in a form of a better performance of bilinguals in tasks tapping into executive function abilities has been reported repeatedly in the literature. However, recent research defends that this advantage does not stem from bilingualism, but from uncontrolled factors or imperfectly matched samples. In this study we explored the potential impact of bilingualism on executive functioning abilities by testing large groups of young adult bilinguals and monolinguals in the tasks that were most extensively used when the advantages were reported. Importantly, the recently identified factors that could be disrupting the between groups comparisons were controlled for, and both groups were matched. We found no differences between groups in their performance. Additional bootstrapping analyses indicated that, when the bilingual advantage appeared, it very often co-occurred with unmatched socio-demographic factors. The evidence presented here indicates that the bilingual advantage might indeed be caused by spurious uncontrolled factors rather than bilingualism per se. Secondly, bilingualism has been argued to potentially affect working memory also. Therefore, we tested the same participants in both a forward and a backward version of a visual and an auditory working memory task. We found no differences between groups in either of the forward versions of the tasks, but bilinguals systematically outperformed monolinguals in the backward conditions. The results are analyzed and interpreted taking into consideration different perspectives in the domain-specificity of the executive functions and working memory.

## The impact of bilingualism on executive functions and working memory in young adults

The core assumption of the bilingual advantage hypothesis (1) is that bilingualism provides enhanced executive function abilities as a consequence of the constant use of two languages. The executive functions (EF) encompass inhibition (i.e., the ability to suppress dominant or salient responses), shifting (the capacity to switch between tasks), and monitoring (the ability to update the information in the working memory; see 2,3). Importantly, as the two languages that a bilingual speaks are always active (4–6), bilinguals require an efficient use of language control: they need to monitor and constantly update the demands of the context they are immersed in and the speakers they are talking to, switch to the target language and inhibit the non-target one (see, for example, the IC model, 7). As it can be seen, language control makes use of EF mechanisms (8). The bilingual advantage hypothesis suggests that active bilingualism trains and improves general EF abilities (see 9–11, for evidence for EF improvement through training) as a consequence of a higher language control demand as compared to a single language use.

The EF have been generally assumed to be domain-general (i.e., the same underlying mechanisms would be responsible for language control and any other kind of executive control, see 12–14). Therefore, if bilingualism trains language control, this training should be then transferred to and captured in any situation that requires the use of domain-general EF abilities. Tasks such as flanker (15), Simon (16) and the Stroop (17) tasks have been classically deemed ideal to explore the bilingual advantage, since all of them tap into general EF abilities (but see below and 18 for a review on little or no convergent validity among tasks). These tasks can provide with indices for different EF components. For example, in all the aforementioned tasks, participants have to deal with congruent trials (trials where all the information presented favors the target response) and incongruent trials (those that present participants conflicting information with the correct response; see Methods section for a more detailed examples of conditions). The difference between the RTs or errors to those two conditions is known as the *Stroop effect* for the Stroop task, and the *conflict effect* for the others, and it’s taken as an indicator of inhibitory abilities. Crucially, those effects have been found smaller for bilinguals when compared to monolinguals in the Stroop task (19) as well as in the Simon (see 20, for young adults; 21, for children; and 1 for seniors) and Flanker task (22). Although it is a view that it has been recanted (23), those findings have been classically interpreted as an evidence for better inhibitory abilities of bilinguals when facing incongruent or conflicting situations, stemming from the constant necessity of inhibiting the non-target language (or stimulus) while managing two languages.

The interpretation of those results, arguing for an improvement in inhibitory abilities for bilinguals, was later challenged by finding advantages in other situations that in principle require no inhibition at all. Thus, Costa et al. (22) found that bilinguals were overall faster than monolinguals in the flanker task (see also 24), not only in the incongruent trials, but also in the congruent ones. As they later argued (25), the improvement in the inhibitory skills would have affected the participants’ responses to the incongruent trials only, not to the congruent trials, where there is nothing to inhibit. Consequently, the authors claimed that bilinguals’ overall faster performance reflected better monitoring abilities, stemming from their constant need of overseeing the linguistic demands of the current environment in order to be able to choose the appropriate language for each communicative situation. This hypothesis gained strength when Costa and his colleagues (25) found that the bilingual advantage in overall reaction times was only found in high-monitoring versions of the tasks (with congruent and incongruent trials evenly distributed) as opposed to low-monitoring conditions (with mostly trials of one kind). In that case, the monitoring advantages in bilinguals seemed noticeable only when the environment was demanding enough (see 1,24, for similar conclusions; but see 18, for a discussion on the impurity of the use of global RTs as a measure of monitoring). Even though these two classic perspectives argued for an advantage in concrete aspects of EF (i.e., monitoring or inhibition), the mixed results prevent researchers to draw strong conclusions as to which EF component was enhanced by bilingualism. Instead, it has been recently argued that a *failure* of bilinguals to inhibit their attention to the non-target language requires a more effortful involvement of EF, causing a more general and unified EF enhancement (23).

The bilingual advantage, however, is not only one of the most popular research topics in the field of bilingualism nowadays, but also one of the most controversial ones (see 26,27, among others, as an example of the growing debate regarding its existence). Some concerns have been raised regarding the results and interpretation of the bilingual advantage (26,28–31). In a nutshell, these authors argue that the bilingual advantage that has been shown in the previous literature is actually a consequence of uncontrolled external factors, small sample sizes and task-dependent effects. Actually, there are many external factors that have been shown to have a direct impact in EF performance, such as participants’ socio-economic status (SES), immigrant status or ethnicity background (32,33). Importantly, these factors tend to differ between bilingual and monolingual populations in certain populations. For example, immigrants –who tend to be bilinguals –show better morbidity and mortality outcomes than non-immigrants around the world (34–39) which is known as “the healthy immigrant” effect. Furthermore, they also display a higher educational profile or IQ level (40–42) than nonimmigrants. Even if it is debatable whether these features are a cause or a consequence of being an immigrant, it seems that individuals who get to pass the immigration screening of host countries display better physical and psychological conditions (43). If those factors are not equal between monolinguals and bilinguals at test, they might potentially cause differences between groups in EF, and there would be no way of disentangling the potential effects of bilingualism from those produced by the uncontrolled factors. Upon reviewing the existing literature showing a bilingual advantage, one could find that the abovementioned concern seems to be the rule more than the exception for the majority of the studies. We observe studies in which SES was not controlled for (13,44), in which comparisons are made between monolinguals and bilinguals that were tested in different countries (45), or in which the bilingual sample tested included the majority of immigrants (19). In addition, and somehow confirming the low reliability and replicability of the bilingual advantage effect, significant findings on bilingual advantage happen principally when sample sizes are small (around n<30, see 31). Furthermore, these effects are not always found across the tasks that are assumed to measure the same construct of executive control (24). As Paap and his colleagues argue, for the hypothesis of the bilingual advantage to be coherently demonstrated, the advantage should be present at least in two different tasks that tap into the same cognitive ability, and the markers of those tasks should correlate, which seems not to be the case (for example, 30). This little or no convergent validity questions the domain-generality of the EF and, as a consequence, the improvement transfer from language control training to enhanced EF provided by bilingualism. Furthermore, when large samples of participants are matched in the confounding variables (26), the bilingual advantage systematically vanishes, with comparable performance or monolingual and bilingual children (46–48), young adults (30) and older adults (49–52). Very recently, Lehtonen et al. (53) explored in detail 891 effect sizes from 152 different studies, both published and unpublished, that compared bilinguals’ and monolinguals’ performance in six different EF tasks. They found very little evidence supporting the bilingual advantage theory, which disappeared when the observed publication bias was corrected for.

Hence, the first aim of the present set of tasks is to test the reliability and replicability of the bilingual advantage in EF. Similarly to what we did with children (46–48) and the very recent work by Dick et al.(54), where they tested more than 4500 children in the flanker, the stop-signal, and the dimensional change card sort tasks, and found little evidence to support the bilingual advantage after conducting Generalized Additive Mixed Models and equivalence test analyses) and the elderly (49), we will test large samples of bilinguals from a purely bilingual community and monolinguals from a monolingual community of the same country, while controlling for potentially confounding factors. The classic tasks used in previous studies reporting bilingual advantage will be used here. To further check for the influence of the controlled factors, bootstrapping analyses will be conducted, exploring whether the significance of a potential bilingual advantage coincides with significant differences in other external factors. Also, the assumption of the domain-generality of the EF will be tested by running correlation analyses between the indices obtained in different tasks (26).

Interestingly, the bilingual advantage has been recently associated with an enhancement of working memory (WM) abilities as well. Similarly to what occurs with EF, it has been shown that WM is also liable to be improved by training (see, among many others, 55–58;but see also 59; for evidence against the beneficial effects of training in WM). However, the relation between bilingualism and WM is a hard domain to explore in isolation, because the concepts of EF and WM are closely connected (60). Indeed, different aspects of EF operate with the information held in the WM, and differences in EF abilities have been argued to correlate with differences in WM capacities (61), especially in complex memory tasks with high demands of storing and processing of information (62). Furthermore, it is important to note that the definitions traditionally given to WM (63) and updating /monitoring (3) overlap in that they both define the ability to manipulate information in the primary memory; and often have been equated (23), although convergent validity analyses have recently shown that they are clearly related but separated components (64). Thus, it has been argued that it is possible that what has been observed as a bilingual advantage in EF might have been a reflection of an advantage in a mediating overlapping factor, i.e. WM. As a consequence, “any finding of an advantage in controlled attention would be far more convincing if WM were held constant” (65). Given that the abovementioned classic EF tasks also require WM capacities, inasmuch as a rule has to be kept in mind to adequately respond to the task, the outcomes would be a product of both EF and WM abilities.

To explore the potential impact of bilingualism on WM abilities, we will also test them in isolation by using the tasks tapping into them alone, where no inhibitory abilities or switching are needed. The studies conducted so far using non-linguistic memory tasks show inconsistent and unclear evidence: Bialystok et al. (1) tested young and old monolinguals and bilinguals in an easy (with two color cues and response possibilities) and a hard (four different cues and responses) version of the Simon task, and found that the increase in difficulty (i.e., increase in the WM load) was handled better by the bilinguals than by the monolinguals (but see 66 for no differences after controlling for socio-demographic factors). Bialystok, Craik, and Luk (19) found no differences between bilinguals and monolinguals in the Self-ordered pointing tasks (67) and minor differences in the Corsi task (68,69), where bilinguals recalled more items than monolinguals, with no differences between forward or backward repetition conditions. Luo, Craik, Moreno and Bialystok (70) tested younger and older monolingual and bilingual adults in forward and backward verbal and spatial (Corsi task) WM tasks, and bilinguals outperformed monolinguals on the spatial tasks but monolinguals did better on verbal tasks. Later, Ratiu and Azuma (71) tested 52 bilinguals and 53 monolinguals in a variety of simple and complex WM tasks, including a backward digit-span task, standard operation span task and a non-verbal symmetry task. Although bilinguals had a significantly higher educational level, which could arguably facilitate the appearance of a bilingual advantage, analyses indicated that monolinguals showed a significantly higher score than bilinguals in operation span and backward digit tasks, with no differences in the symmetry task. However, in a multivariate regression analysis, speaker group (bilinguals vs. monolinguals) did not predict scoring in any of the tasks, nor it interacted with any other factor. Interestingly, we observe that some of the concerns with respect to the influence of external factors (26,30,31) do apply to these studies also. Namely, the bilinguals tested by Bialystok, Craik, and Luk (19) spoke a huge variety of second languages, indicating different cultural (and probably ethnical) backgrounds, and the SES was not measured. Crucially, out of 24 participants per group, 14 young bilinguals and 20 old bilinguals were immigrants. In the study by Luo, Craik, Moreno and Bialystok (70), language groups differed in their English vocabulary level and the nonverbal intelligence scores. This was to some extent solved by including those factors in an ANCOVA showing that the effect remained the same, but still their samples feature uneven sizes (58 monolinguals and 99 bilinguals) and language backgrounds (the second language spoken by the young adults varied among a set of more than 10 languages), suggesting differences in ethnicity. Importantly, many other relevant variables such as immigrant status or SES were not reported.

Recently, trying to account for WM differences while controlling for external factors, Hansen and colleagues (72) tested 152 native Spanish children, half of whom were attending a bilingual immersion schooling program (i.e., receiving teaching in both English and Spanish) and the other half Spanish only. Both groups of children were matched in various demographical factors, including SES and intelligence, and they all were from the same city. They were tested in an n-back task (participants have to indicate if the current trial matches the one from n-trials back) and a reading span task (a task that requires remembering the last word of a presented set of sentences), as well as a rapid automatic naming task to measure participants’ verbal processing speed. They found that young bilinguals performed better than monolinguals in the n-back tasks, a task that heavily relies on updating (monitoring and updating the items held in the WM) and inhibition (the relevant item changes in every trial). On the other hand, young bilinguals showed a disadvantage in the reading span task (which requires mostly linguistic processing and verbal storage), but an advantage in older groups (see also 73, for an extended exploration on the effects of emergent bilingualism on literacy). Hansen and colleagues argue for an effect of bilingualism on WM during the first years of immersion due to the higher demands that children might face when they encounter bilingualism for the first time. But, this difference would just modulate the development of said skills, and importantly they would equalize eventually (on the same regard, see also 74). However, in a similar longitudinal perspective to explore the effect of bilingualism in WM during adulthood, Ljunberg and colleagues (75), found that bilinguals systematically outperformed monolinguals ranging from 35 to 80 years of age in several tasks tapping into episodic memory and letter fluency tests.

As it can be seen, there are few studies that specifically looked at the effects of bilingualism on WM. The pattern of results ranges from similar performance of bilinguals and monolinguals (24,76) to benefit of bilingualism (19,74,75), or even bilingual disadvantage (71). Considering the opposing pieces of evidence presented so far and the lack of control of the relevant factors (26,30,31) that some studies feature, consistent conclusions cannot be drawn from those data. Therefore, the effects of bilingualism on WM should be tested with the samples that only differ in linguistic background, with bilingual and monolingual profiles as similar as possible.

The current experiment aims at testing the effects of bilingualisms on both EF (first set of four tasks) and WM (second set of four tasks) with large cohorts of carefully matched monolinguals and bilinguals. Ninety young bilingual adults from the Basque Country (a region of the north of Spain where Basque and Spanish are co-official) and 90 carefully matched monolinguals from Murcia (a south-eastern region of Spain where only Spanish is spoken and official) were tested. To explore the assumption of the bilingual advantage theory that claims that bilingualism enhances general EF that applies to any domain general situation, the degree of cross-task replicability will be also tested (30). Also, to account for the possible impact of bilingualism on WM, the forward and backward versions of a spatial (Corsi task) and a numerical (digit span task) memory tasks will be used. Thus, while it is impossible to get rid of any possible WM influence in the set of EF tasks, the use of more specific tasks will provide us with information about WM in isolation. To assess the co-occurrence of a bilingual advantage and significant socio-demographic differences, additional bootstrapping and regression analyses will be also conducted. In sum, we tested the possible benefits of bilingualism on two different aspects, i.e., EF and WM, in large samples of bilinguals and monolinguals matched on all the relevant factors.

### General Methods

#### Participants

180 young adults from Spain took part on these series of tasks. The 90 bilinguals (68 females, mean age 22.29 year, SD= 2.87) were tested in the facilities of the BCBL, in Donostia-San Sebastian (in the Basque Country). On average they had acquired Basque with 0.96 years of age (SD=1.27) and they reported to have a general proficiency of 8.41 over 10 (SD=1.88) in Basque. Self-reports in Spanish proficiency reached a mean score of 8.58 (SD=1.91), and this language was acquired with an average of 1.13 years (SD=1.72). Thus, bilinguals were balanced in terms of proficiency (*p*>.33) and age of acquisition (*p*>.42). They were also interviewed by native bilingual research assistant to make sure that they were truly balanced and native bilinguals. As opposed to heritage bilingual speakers, Basque bilinguals tend to show a more balanced dominance and exposition to both languages, as they are fully immersed in a bilingual society. Generally, Spanish is more present in the media and leisure activities, but the bilinguals from our sample report to be equally exposed to Spanish (47.78% of their time, SD=17.07) and to Basque (43% of their time, SD= 17.95), with no statistical differences (*p*>.18). The bilingual educational system allows for tuition models based on either one of the languages or both, and although data on this was not collected, self-reported percentage of time in which they write in each of the languages – which could be taken as a proxy of the language in which they carry over their educational activity – was collected. We found out that they write more in Basque (52.67% of their time, SD=24.30) than in Spanish (39%, SD=23.80, *p*<.01), indicating a predominant tendency for formal education in Basque. On average, they report that they speak 50% of the time in Spanish (SD=19.11) and 43.22% in Basque (SD=20.21), a different that although results in marginally significant (*p*>.08), reflects a highly balanced use of both languages.

The 90 monolinguals (67 females, 21.84 years of age in average, SD=3.05) were recruited in the region of Murcia, in the south-east area of Spain, and tested in the University of Murcia. They reported to have acquired Spanish with a mean age of 0.68 (SD=.76) and a mean proficiency of 9.13 (SD=.84), with very little or no knowledge of any other language.

Participants from both groups were matched for a variety of factors that could potentially affect our experimental purposes. The matched 90 bilinguals and 90 monolinguals were selected from a bigger sample of 126 monolinguals and 141 bilinguals by means of testing their differences in age, IQ, socio-economic status (SES), educational level and knowledge of Spanish using independent sample t-tests. An estimation of the IQ of each participant was based on their performance on an abridged version of the Kaufman Brief Intelligence Test (K-BIT, 77) that was administrated during the experimental session. As an indicator of the SES, total monthly income was considered and divided by the amount of household members, thus getting an approximate value of the incomes that each member of household receive monthly on average and make the incomes more comparable across families of different sizes. The majority of the participants had already obtained a university degree (or higher) or were in the process of obtaining one, and this number was virtually identical across groups (88 bilinguals and 87 monolinguals). To control for their proficiency in Spanish (namely, the test language) every participant completed the Spanish version of the LexTale (78) that provides an objective indicator of their Spanish mastery. All these demographic and linguistic variables that could affect the outcomes of the study were thus matched across groups (all *p*s>.1; see Table 1 for detailed information about the participants). All participants reported normal or corrected to normal vision and signed a consent form according to the principles established by the ethics committee of the BCBL.

**Table 1:** Demographic factors of the participants. Mean values are presented together with standard deviation (between parentheses) and the p value resulting from an independent groups t-test with an alpha value of 0.05.

|  | Monolinguals |  | Bilinguals |  | p value |
| --- | --- | --- | --- | --- | --- |
| Chronological age (years) | 21.84 | (3.05) | 22.3 | (2.87) | 0.31 |
| General IQ | 22.76 | (2.62) | 23.4 | (2.91) | 0.13 |
| SES (income in €/household members) | 639.55 | (498.97) | 739.58 | (297.36) | 0.1 |
| LexTale | 92.28 | (5.63) | 93.4 | (3.88) | 0.11 |

#### Procedure

Bilingual participants were tested in the facilities of the BCBL in Donostia-San Sebastián, and monolingual participants were tested in University of Murcia, in Murcia. In both locations, participants went through the experimental session in a room with equivalent settings, with the same equipment used in both labs. The experiment was run using Experiment Builder (© SR Research) v 1.10.1385, and the CRT monitor was set to 60Hz in a resolution of 1280×1024 and placed at an approximate distance of 60 centimeters from the participants. Manual responses were recorded using a response box with 7 buttons, being the first one on the left painted in red and the last one on the right painted in green. When audio was played or voice inputs were recorded, it was done by using Sennheisser PC151 headsets.

Following four pseudo-randomized lists, which were the same for both bilinguals and monolinguals, participants performed four tasks aimed at measuring their EF and four other tasks aimed at gathering information about their WM skills. The tasks used to measure EF were the Flanker task (15),Simon task (16), the Verbal Stroop task (17), and the Numerical Stroop task (79). The tasks used to measure WM were two versions (forward and backward) of the Corsi test (67,68) and the digit span test (80). All the tasks were conducted in the same experimental session in one day, which lasted around 60 minutes. The duration was approximate and contingent on the participants’ performance (see below for details). The results of the tasks were analyzed following the traditional alternative hypothesis testing with ANOVAs, as well as using Bayesian Null Hypothesis testing comparisons (81,82).

### Executive Functions: Tasks 1-4

#### Materials and procedure

For the <u>flanker task</u>, rows of five arrows (←) were displayed on the center of the screen. For the *congruent* condition, the central arrow was flanked by four arrows pointing in the same direction (← ← ← ← ←). For the *incongruent* condition, the central arrow was flanked by arrows pointing in the opposite direction (← ← → ← ←), and for the *neutral* condition, the arrow was flanked by no arrows (-- -- ← -- --). There were 16 items of each condition, 8 of them with the central arrow pointing to the left and the other half with the central arrow pointing to the right.

In the <u>Simon task</u> participants were presented with a black circle or a black square in the screen, and they were instructed to respond with the red button (on the left of the response box) if they saw a circle or with the green button (on the right) if they saw a square, irrespectively of its position in the screen. Thus, the *incongruent* condition was created by presenting circles on the right side of the screen or squares presented on the left side of the screen, making participants respond to them with the button on the opposite side of the side in which the figure appeared. The *congruent* condition was created by presenting the figures in the same side of the screen of the response button needed, i.e., by presenting circles in the left and squares in the right. Finally, the *neutral* condition was created by presenting the figures in the middle of the screen. There were 16 items of each condition, and half of the items per condition were squares, while the other half were circles.

For the <u>verbal Stroop</u> task, the Spanish words for the colors red, blue and yellow (“rojo”, “azul” and “amarillo”) and three pairwise-matched (with a similar length, frequency and syllabic structure) non-color words (“ropa”, “avión” and “apellido”, the Spanish words for clothes, plane and surname, respectively) were used as target items. They were arranged to create the *congruent* (a color word printed in the same color that the word indicates; e.g., the word “azul” in blue), *incongruent* (a color word printed in a different color from what it is naming, e.g., the word “rojo” in blue) and *neutral* (non-color words printed in any of the colors, e.g., the word “ropa” in red) conditions. In each condition 24 trials were used, and each color was presented the same amount of times in each condition (8 times), paired equally with every word. All the strings were presented in uppercase Courier New font on a black background, while the colors were set in the RGB-scale values as follows: blue=0,0,255; red=255,0,0; yellow=255,255,0.

For the <u>numerical Stroop</u> task, stimuli consisted in six digits (2, 3, 4, 6, 7, and 8), arranged in pairs to form each trials (e.g., 2-6), one presented on the left side of the screen and another one on the right side. Participants had to say which one was larger in size, ignoring the numerical value. Depending on how the digits were paired, three conditions were created: 24 *congruent* trials (the larger number in magnitude was also the bigger in size, e.g., small 2-big 6), 24 *incongruent* trials (the smaller number in size was the larger in magnitude, e.g., big 2-small 6) and 24 *neutral* trials (two same numbers different in size, e.g., big 4-small 4). In all the conditions “left” and “right” responses were equally distributed, and each digit was used in each condition an equal number of times.

Participants were instructed about the responses before each task, and received indications about the button press or vocal responses in each case. In the Simon, the Flanker, and the Numerical Stroop tasks, there was a short training phase before the experiment started. After a fixation point was displayed in the center of the screen for 1000ms in black, on a white background, the stimuli was presented on the screen for 5000ms or until the response was given. The order of the stimuli was randomized and there were no breaks. In the verbal Stroop task, after a short training period, a fixation mark was presented for 250ms (a white cross in a black background), and then the target word appeared on the screen for 3000ms, during which participants’ response was recorded.

#### Data analysis

The data from the <u>Stroop task</u> needed a different pre-processing before analyzing. Audios were equalized to a 63dB amplitude using Praat© (83). Once all the files had same amplitude level, the voice onset was automatically detected by Praat as follows: the textgrid of each audio was divided into “sound” and “silence” segments using the silence function from Praat. For a segment to be considered “sound” it had to have a minimum pitch of 100 Hz, to have exceeded a −25dB threshold and to have lasted at least 100ms. “Silence” segments had to last at least 200ms. The starting time point of the first sound segment was considered the onset of the speech and therefore, the reaction time of that response. The accuracy of the responses was checked manually, and the speech onset was manually adapted in the cases in which subjects corrected themselves (e.g., “roj…amarillo”) and mistakes were removed.

For the four tasks, after removing errors, latencies were trimmed for outliers by deleting any response that deviated in more than 2SD from the mean in each condition. After this, a 3×2 ANOVA was run with Condition (*congruent, incongruent, neutral*) and Language Group (*bilinguals, monolinguals*). To further check for any possible advantage, the *conflict index* (i.e., *incongruent-congruent* latencies), the *incongruity index* (i.e., the *neutral* condition compared to the *incongruent* condition), and the *congruency index* (the *congruent* condition compared to the *neutral* one) were compared across language groups using ANOVAs and Bayesian t-tests (calculated using JASP,84). The same procedure was repeated for error rates.

#### Results

In the reaction times analysis for the <u>flanker task,</u> after removing 5.15% of the data due to outliers, we observed a strong main effect of Condition [*F*(2, 356)=196.15, *p*<.01], and a more detailed analysis indicated that *congruent* items were responded faster than the *incongruent* ones [*t*(179)= 16. 69, *p*<.01], and also *neutral* items were responded faster than both *incongruent* [*t*(179)= 15.98, *p*<.01] and *congruent* ones [*t*(179)= 2.48, *p*<.02]. There was no main effect of Language Group [*F*(1, 178)=0.60, *p*>.44] and no interaction between the two main effects [*F*(2, 356)= 0.01, *p*>0.99]. The *conflict index* analysis showed a strong Condition effect [*F*(1,178)=279.92, *p*<.01], but no other effect or interaction were significant (*F*s<1). Results from the Bayes Factor t-test comparison across groups [*BF*_01_=6.14] indicated that the null hypothesis explains the data 6 times better than the alternative hypothesis of bilinguals showing a reduced *conflict index* (see Table 2 for descriptive results). The *incongruity index* showed the same pattern as the *conflict index*, showing a strong Condition effect [*F*(1, 178)= 253.96, *p*<.01] but no other significant results (*F*s<1), and the Bayesian Factor analysis favored the null hypothesis [*BF*_01_=5.16].

**Table 2:** Flanker task. Mean reaction times (in milliseconds) and error rates (in percentages) for each condition and index are displayed together with standard deviations (between parentheses).

|  |  | Mean reaction times |  |  |  | Mean error rates |  |  |  |
| --- | --- | --- | --- | --- | --- | --- | --- | --- | --- |
|  |  | Monolinguals |  | Bilinguals |  | Monolinguals |  | Bilinguals |  |
| Conditions | Congruent | 387 | (67) | 379 | (80) | 0.49 | (1.68) | 0.49 | (1.68) |
|  | Incongruent | 428 | (78) | 420 | (89) | 2.71 | (4.4) | 2.85 | (4.97) |
|  | Neutral | 382 | (64) | 373 | (68) | 0.63 | (1.89) | 0.69 | (2.19) |
|  | Total | 399 | (67) | 391 | (77) | 1.27 | (1.97) | 1.34 | (2.03) |
| Effects | Stroop | 41 | (33) | 41 | (34) | 2.22 | (4.33) | 2.36 | (4.92) |
|  | Incongruity | -46 | (38) | -47 | (41) | -2.08 | (4.29) | -2.15 | (5.48) |
|  | Congruency | 6 | (29) | 6 | (32) | 0.14 | (2.29) | 0.21 | (2.38) |

The *congruency index* also showed a significant effect of Condition [*F*(1, 178)= 6.14, *p*<.02] but no other significant effect or interaction (*F*s<1). Bayesian analysis also favored the null hypothesis over the alternative one [*BF*_01_=4.71].

The analysis of the error rates showed a similar pattern. A strong and significant Condition effect was found [*F*(2, 356)=34.39, *p*<.01], stemming from *incongruent* trials producing more errors than the ones belonging to *congruent* condition [*t*(179)= 6.66, *p*<.01] and to *neutral* trials [*t*(179)= 5.79, *p*<.01]; but no differences were found when *congruent* and *neutral* conditions were compared [*t*(179)= 1.00, *p*>.32]. Importantly, no main effect of Language Group or an interaction between it and Condition were found (all *Fs*<1). When the *conflict index* was computed and compared between groups, the effect of Condition was significant [*F*(1,178)=44.06, *p*<.01], but no other main effect or modulation was (*F*s<1). Expectedly, the Bayes Factor t-test analysis [*BF*_01_=6,07] supported the null-differences hypothesis. Similarly to what it was found in the RTs analysis, the *incongruity index* was significant [*F*(1, 178)= 33.35, *p*<.01] but no other effect or interaction was (*F*s<1), as confirmed by the Bayesian Factor analysis [*BF*_01_=6.16]. *Congruency index* analysis, however, showed no main effect or interaction (all *F*s<1), but still the index was compared across groups and the null hypothesis was the best fir for the data [*BF*_01_=6.08].

The general ANOVA conducted on the reaction times to correct responses –after removing the outliers (4.82% of the data) – in <u>the Simon task</u> revealed a significant main effect of Condition [*F*(2, 356)=28.66, *p*<.01]. Post-hoc comparisons showed that *incongruent* trials were responded slower than both *congruent* [*t*(179)= 8.09, *p*<.01] and *neutral* trials [*t*(179)= 4.71, *p*<.01], and that *congruent* trials were responded faster than *neutral* ones [*t*(179)= 2.38, *p*<.02]. However, nor main effect of Language Group [*F*(1, 178)=1.91, *p*>.17] neither the interaction between them [*F*(2, 356)=0.33, *p*>.72] was significant (see Table 3 for descriptive results). In the *conflict index* analysis, Condition effect was significant [*F*(1, 178)= 65.01, *p*<.01], but no main effect of Language Group [*F*(1, 178)= 1.64, *p*>.2] or an interaction was found [*F*<1], which indicates that there are no significant differences in the *conflict index* when it is compared across groups [*BF*_01_=5.90]. The *incongruity index* showed a strong Condition effect [*F*(1, 178)= 22.01, *p*<.01], but Language Group did not [*F*(1,178)= 1.89, *p*>.17] and neither did it the interaction between them [*F*<1]. The Bayes Factor analysis supported that the *incongruity index* was similar in both groups [*BF*_01_=4.73]. In a similar pattern, the *congruency index* was significant [*F*(1, 178)= 5.66, *p*<.02] but it did not interact with Language Group [*F*<1]. Language Group did not result significant either [*F*(1, 178)= 2.09, *p*>.15], and Bayes Factor analysis supported the similarity of the index across language groups [*BF*_01_=5.48].

**Table 3:**
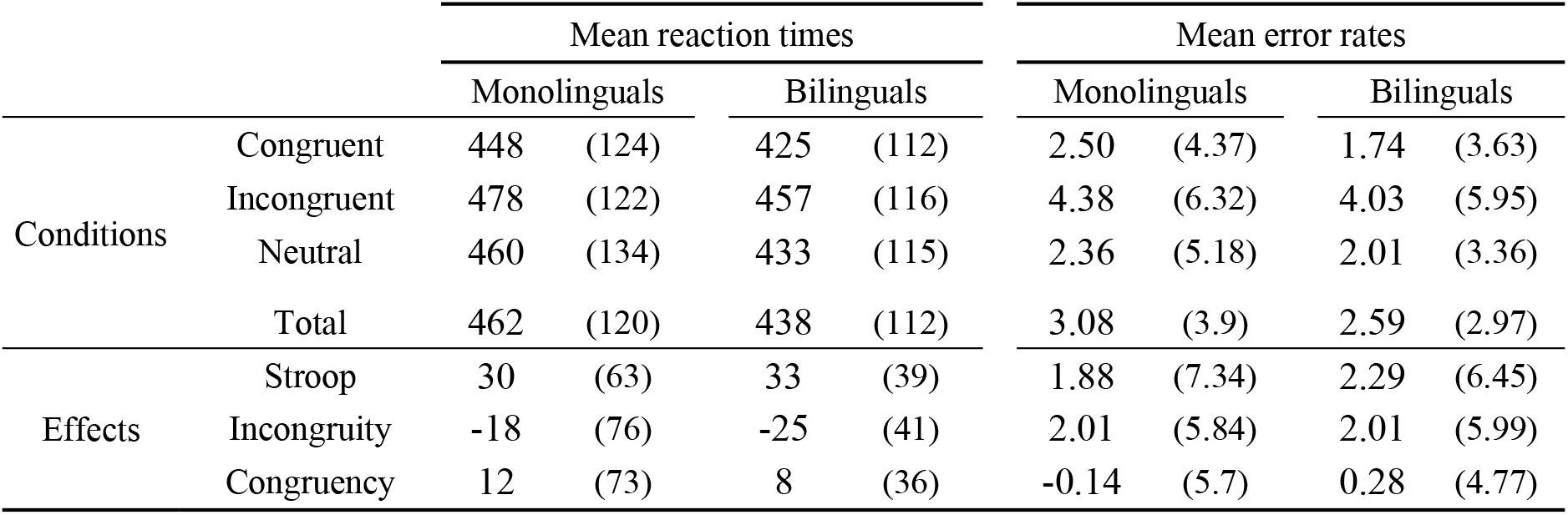
Simon task. Mean reaction times (in milliseconds) and error rates (in percentages) for each condition and index are displayed together with standard deviations (between parentheses).

When the error rates were analyzed, a similar picture emerged. Condition was significant [*F*(2, 356)=13.7, *p*<.01], and paired comparisons revealed that it was due to the *incongruent* trials producing more errors than both *congruent* [*t*(179)= 4.05, *p*<.01] and *neutral* ones [*t*(179)= 4.58, *p*<.01], with no difference between these last two (*t*<1). No effect of Language Group or an interaction between it and Condition were found (all *Fs*<1). Crucially, the *conflict index* was significant [*F*(1,178)= 16.35, *p*<.01] but it did not interact with Language Group, and Language Group did not result significant either (all *F*s<1). The Bayes Factor analysis [*BF*_01_=5.74] indicated that the null-hypothesis was almost 6 times more likely to explain the data. Similarly, the *incongruity effect* was significant [*F*(1,178)= 20.88, *p*<.01] but neither language Group nor the interaction between it and Condition was (all *F*s<1). The Bayes Factor analysis indicated that the index was highly similar across language groups [*BF*_01_=5.79]. The analysis of the *congruity effect* revealed that neither Condition, nor Language group nor the interaction between them was significant (all *F*s<1.3, all *p*s>.26).

After removing the outliers from the <u>Stroop task</u> (4.84%), a general ANOVA on the reaction times to correct responses showed a main effect of Condition [*F*(2, 356)=279.22, *p*<.01], which showed that *congruent* condition was responded on average faster than *neutral* trials [*t*(179)= 10.98, *p*<.01] and than *incongruent* trials [*t*(179)= 21.32, *p*<.01]. *Neutral* condition was also responded faster than *incongruent* condition [*t*(179)= 13.80, *p*<.01]. Crucially, we observed no main effect of Language Group [*F*(1, 178)=1.53, *p*>.22] and no interaction between it and Condition [*F*(2, 356)=0.40, *p*>0.67]. The *Stroop index* analysis indicated a strong effect of Condition [*F*(1, 178)= 452.41, *p*<.01], but Language Group was not significant [*F*(1, 178)=1.3, *p*>.29] and, crucially, it did not interact with Condition (*F*<1). Importantly, the analysis of the Bayes Factor [*BF*_01_=5.62] indicated that there are no significant differences between groups and that the null hypothesis is the most likely one to explain these data (see Table 4 for descriptive results). The *incongruity index* analysis showed a main effect of Condition [*F*(1,178)= 189.59, *p*<.01] but negligible main effect of Language Group [*F*(1,178)= 1.77, *p*>.18] and, importantly, no modulation of Condition by Language Group (*F*<1). This null difference between groups was again supported by the Bayesian t-test [*BF*_01_=5.79]. The analysis of the *congruency index* showed a significant effect of Condition [*F*(1,178)= 120.32, *p*<.01] but it was not modulated by the knowledge of a second language [*F*(1,178)= 1.06, *p*<.30], which was supported by the Bayesian t-test [*BF*_01_=3.78]. Similarly, no main effect of Language Group was found [*F*(1,178)= 1.62, *p*>.21].

**Table 4:** Verbal Stroop task. Mean reaction times (in milliseconds) and error rates (in percentages) for each condition and index are displayed together with standard deviations (between parentheses).

|  |  | Mean reaction times |  |  |  | Mean error rates |  |  |  |
| --- | --- | --- | --- | --- | --- | --- | --- | --- | --- |
|  |  | Monolinguals |  | Bilinguals |  | Monolinguals |  | Bilinguals |  |
| Conditions | Congruent | 648 | (96) | 662 | (96) | 0.05 | (0.44) | 0.05 | (0.44) |
|  | Incongruent | 743 | (116) | 761 | (104) | 0.65 | (1.97) | 0.32 | (1.43) |
|  | Neutral | 684 | (86) | 705 | (98) | 0.14 | (0.75) | 0.14 | (0.75) |
|  | Total | 692 | (94) | 709 | (95) | 0.28 | (0.84) | 0.17 | (0.72) |
| Effects | Stroop | 95 | (65) | 99 | (58) | 0.60 | (1.93) | 0.28 | (1.37) |
|  | Incongruity | -60 | (63) | -56 | (50) | -0.51 | (1.75) | -0.19 | (1.24) |
|  | Congruency | 35 | (48) | 43 | (47) | 0.09 | (0.88) | 0.09 | (0.62) |

The error rate analysis showed a strong and significant Condition effect [*F*(2, 356)=10.24, *p*<.01], indicating that more errors were made in the items belonging to the *incongruent* condition than in the ones belonging to the *congruent* condition [*t*(179)= 3.52, *p*<.01] and to the *neutral* one [t(179)= 3.07, *p*<.01]; but no effect of Language Group was found [*F*(1,178)=0.87, *p*>.35] nor an interaction between Language Group and Condition [*F*(2, 356)=1.67, *p*>0.19]. The *Stroop index* was computed and compared between groups, which showed a strong main effect of Condition [*F*(1,178)= 12.43, *p*<.01], but it was not modulated by Language Group [*F*(1,178)= 1.69, *p*>.2], and Language Groups did not differ either [*F*(1,178)= 1.35, *p*>.25]. Furthermore, the Bayes Factor analysis [*BF*_01_=2.83] supported the null-differences hypothesis. The *incongruity index* was significant [*F*(1,178)= 9.45, *p*<.01], but Language Group did not modulate it [*F*(1,178)= 2.06, *p*>.15] and neither a main effect of Language Group was observed (*F*<1). The Bayes factor comparison also tended to support the null hypothesis as the best fitting candidate [*BF*_01_=2.38]. In the *congruency index* analysis, Condition was not significant [*F*(1,175)=2.67, *p*>.1] and neither was the effect of Language Group or the interaction between the two factors (*F*s<1).

In <u>numerical Stroop task</u>, the reaction times to the correct responses were analysed after removing of outliers (4.75%). The general ANOVA showed a main effect of Condition [*F*(2, 356)=202.38, *p*<.01], indicating that *congruent* trials were responded faster than both the *incongruent* [*t*(179)= 16.37, *p*<.01] and the *neutral* ones [*t*(179)= 6.04, *p*<.01], and that *neutral* items were also responded faster than the *incongruent* ones [*t*(179)= 13.80, *p*<.01]. Crucially we found no main effect of Language Group [*F*(1, 178)=2.61, *p*>.11] nor interaction between it and Condition [*F*(2, 356)=0.40, *p*>0.67]. The *Stroop index* was significant [*F*(1,178)= 268.63, *p*<.01] but Language Group [*F*(1,178)= 2.95, *p*>.09] was not. The lack of interaction between them [*F*(1,178)=1.44, *p*>.23] indicated that the linguistic profile did not have any reliable impact on the magnitude of the *Stroop effect* [*BF*_01_=3.18] (see Table 5 for descriptive results). The *incongruity index* analysis showed a significant Condition effect [*F*(1,178)= 191.70, *p*<.01], but Language Group was not significant [*F*(1,178)= 2.62, *p*>.11] and neither it was the interaction between them [*F*(1,178)= 2.23, *p*>.14], which was supported by the tendency showed by the Bayes Factor analysis [*BF*_01_=2.2]. The *congruency index* was strong [*F*(1,178)=32.25, *p*<.01], but Language effect was not significant [*F*(1,178)=2.19, *p*>.14], neither was the interaction between them (*F*<1). The null hypothesis was also supported by the Bayes Factor analysis [*BF*_01_=5.89].

**Table 5:** Numerical Stroop task. Mean reaction times (in milliseconds) and error rates (in percentages) for each condition and index are displayed together with standard deviations (between parentheses).

|  |  | Mean reaction times |  |  |  | Mean error rates |  |  |  |
| --- | --- | --- | --- | --- | --- | --- | --- | --- | --- |
|  |  | Monolinguals |  | Bilinguals |  | Monolinguals |  | Bilinguals |  |
| Conditions | Congruent | 424 | (86) | 404 | (86) | 0.37 | (1.35) | 0.37 | (1.35) |
|  | Incongruent | 474 | (103) | 447 | (100) | 2.27 | (4.29) | 2.69 | (4.43) |
|  | Neutral | 433 | (95) | 414 | (91) | 0.23 | (1.15) | 0.32 | (1.56) |
|  | Total | 444 | (93) | 422 | (90) | 0.96 | (1.75) | 1.13 | (1.68) |
| Effects | Stroop | 51 | (36) | 44 | (41) | 1.90 | (4.42) | 2.31 | (4.47) |
|  | Incongruity | -41 | (37) | -33 | (35) | -2.04 | (3.81) | -2.36 | (4.72) |
|  | Congruency | 10 | (25) | 11 | (20) | -0.14 | (1.7) | -0.05 | (2.02) |

The error rate analysis also showed a significant Condition effect [*F*(2, 356)=38.79, *p*<.01], which was a reflection of *incongruent* trials producing more errors than both *congruent* [*t*(179)= 6.26, *p*<.01] and *neutral* [*t*(179)= 6.79, *p*<.01] ones, but no differences were found between *congruent* and *neutral* items (*t*<1). Importantly, we observed no effect of Language Group or an interaction between that factor and Condition (all *Fs*<1). The *Stroop index* was compared between groups, and it revealed a main effect of Condition [*F*(1,178)= 40.43, *p*<.01], but no other effects were significant (all *F*s<1). Bayes Factor analysis [*BF*_01_=4.80] indicated that both groups behaved similarly. Similarly, the *incongruity effect* analysis showed a Condition effect [*F*(1,178)= 47.32, *p*<.01], but no main effect of Language or interaction was found (all *F*s<1). Again, Bayes Factor analysis indicated no differences between Language Groups [*BF*_01_=5.49]. Finally, the *congruency index* analysis showed no significant effect or interaction (all *F*s<1), and a further Bayes Factor analysis also indicated that the index did not differ across groups [*BF*_01_=5.88].

### Cross-task correlation

The cross-task coherence was tested in a correlation analysis between the indices in milliseconds (*congruency, incongruity* and *conflict/Stroop*) obtained in all the tasks (30,64). The Stroop/conflict effects across tasks showed negligible correlation strength (all rs between −.06 and .10). The *congruency* effect showed a significant yet low positive correlation between the flanker task and the numerical Stroop task (r= .21), but all the other effects reflected mild cross-task reliability (all rs between −.05 and .15). Finally, analyses on *incongruity effects* showed a significant but negative correlation between the flanker and the Simon tasks (r= −.20,) and all the rest pairs of effects indicated that the cross-task coherence was very low (all rs between −.11 and .17). Thus, this analysis consistently showed a poor level of cross-task correlation.

### Interim conclusion

We found no evidence that supports a bilingual advantage in EF: bilinguals and monolinguals performed equally when facing incongruent trials or when measuring global reaction times. Furthermore, the much more restrictive Bayesian Factor analysis of the different *Stroop* and conflict effects clearly favored the null hypothesis, reinforcing the idea of bilinguals and monolinguals performing similarly in classic tasks that tap into EF. The indices obtained in the different tasks showed negligible or, when significant, negative correlation between them.

As explained in the Introduction, in the second part of the present study we aimed at exploring the potential differences between groups in WM. With that purpose, we tested the same participants in two tasks that measure WM –a verbal/numerical WM task and a spatial WM task – in both a forward and a backward version.

### Working memory: Tasks 5-8

#### Materials and procedure

In the <u>Corsi</u> and the <u>Corsi inverse</u> tasks, the participants were presented with 10 blue squares distributed on a screen with a grey background (see Fig 1 for the distribution of the squares in the screen). Those blue squares would change into green in previously established orders, creating the sequences that the participants had to remember. The ten squares were firstly presented in the screen, in blue, for 1000ms. After that, one of them changed to green for 1000ms. Then they all went back to blue for another 1000ms, and the next square in the sequence changed to green. This process was repeated until all the squares of the current sequence were turned to green and back to blue again. Then, a question mark appeared in the middle of the screen, and participants needed to point with the finger which squares changed their color. In the <u>Corsi</u> task, participants had to indicate the changes in the same order as they occurred. In the <u>Corsi inverse</u> task, they had to indicate the changes in the reverse order as they occurred. There were 8 consecutive blocks with an increasing difficulty level, and in each block two sequences (trials) were presented. The increase in the difficulty was produced by the addition of one more square change to the sequences of each consecutive block. Thus, the trials in the first block consisted of two square changes, the trials in the second block consisted of three square changes, and so on up to 9 square changes in the last block. After each block (that included 2 trials), the experimenter noted the number of correctly recalled sequences. If the participant failed both, the experiment ended. If the participant remembered one or two, the next block started (see Fig 2 for a schematic representation of the experiment, and see Table 6 for the details on each trial).

**Fig 1:**
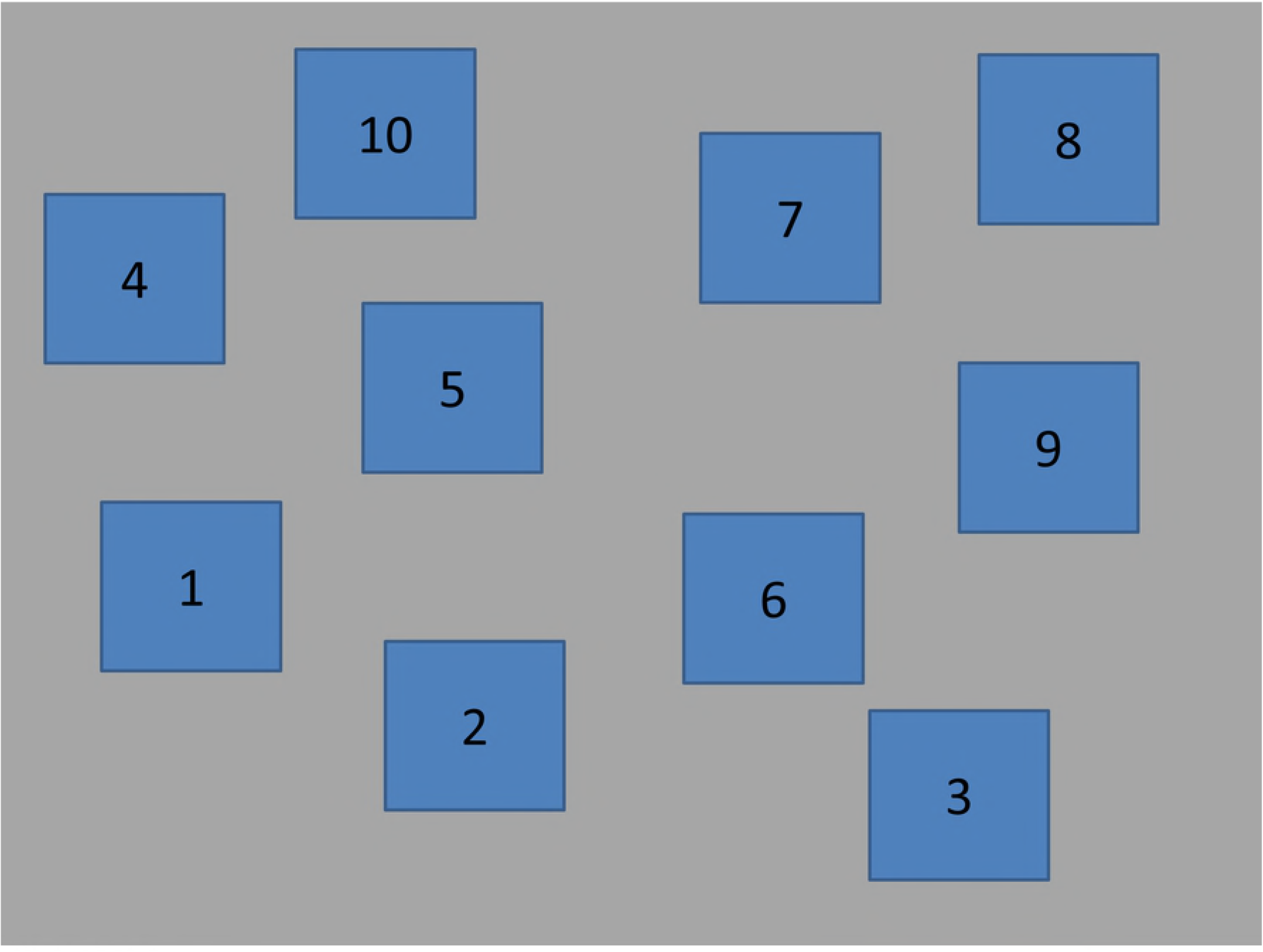
Spatial distribution of the squares in the Corsi and Corsi inverse tasks and the numbers assigned to each of them.

**Fig 2:**
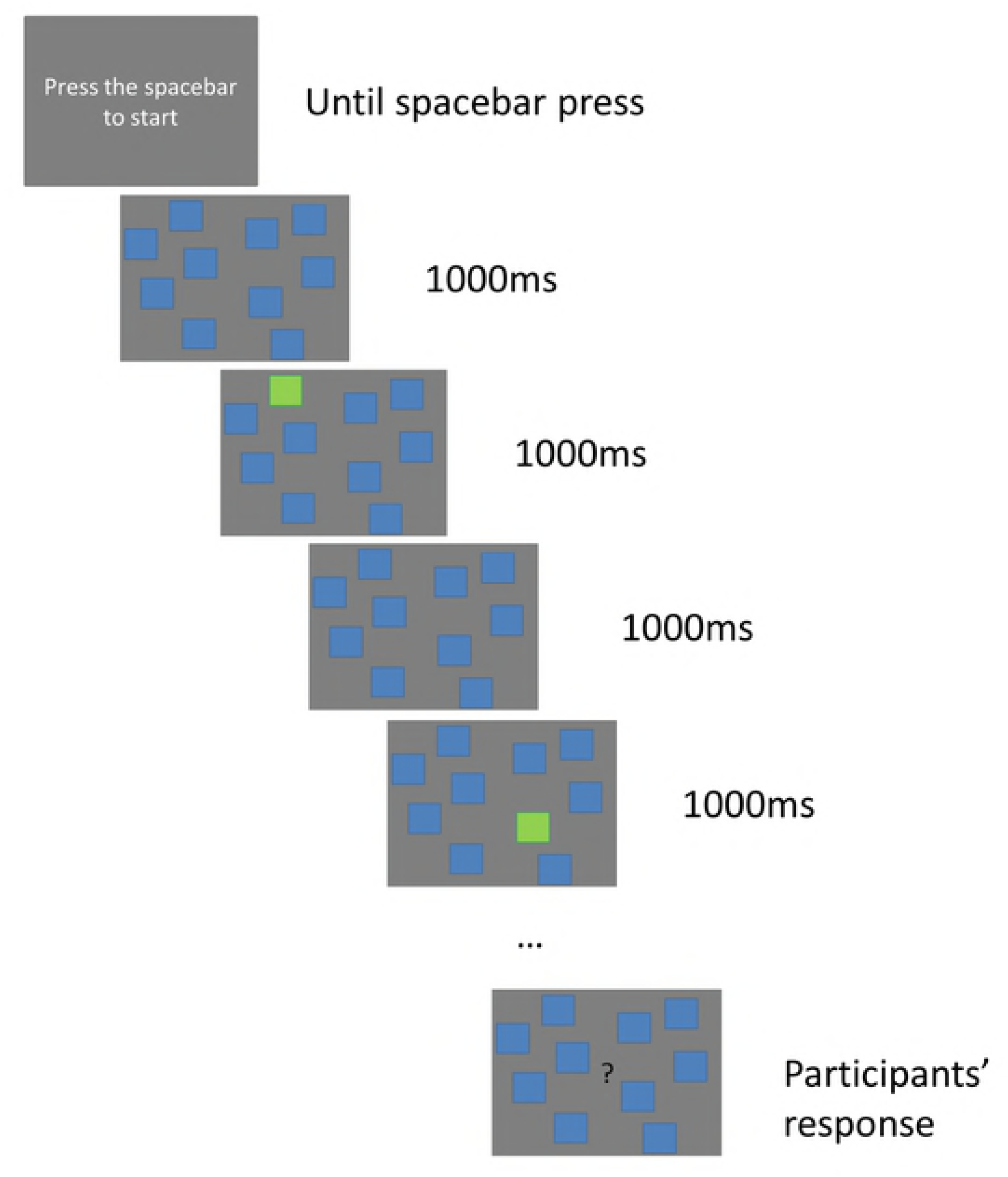
Schematic representation of the Corsi and Corsi inverse tasks.

**Table 6:**
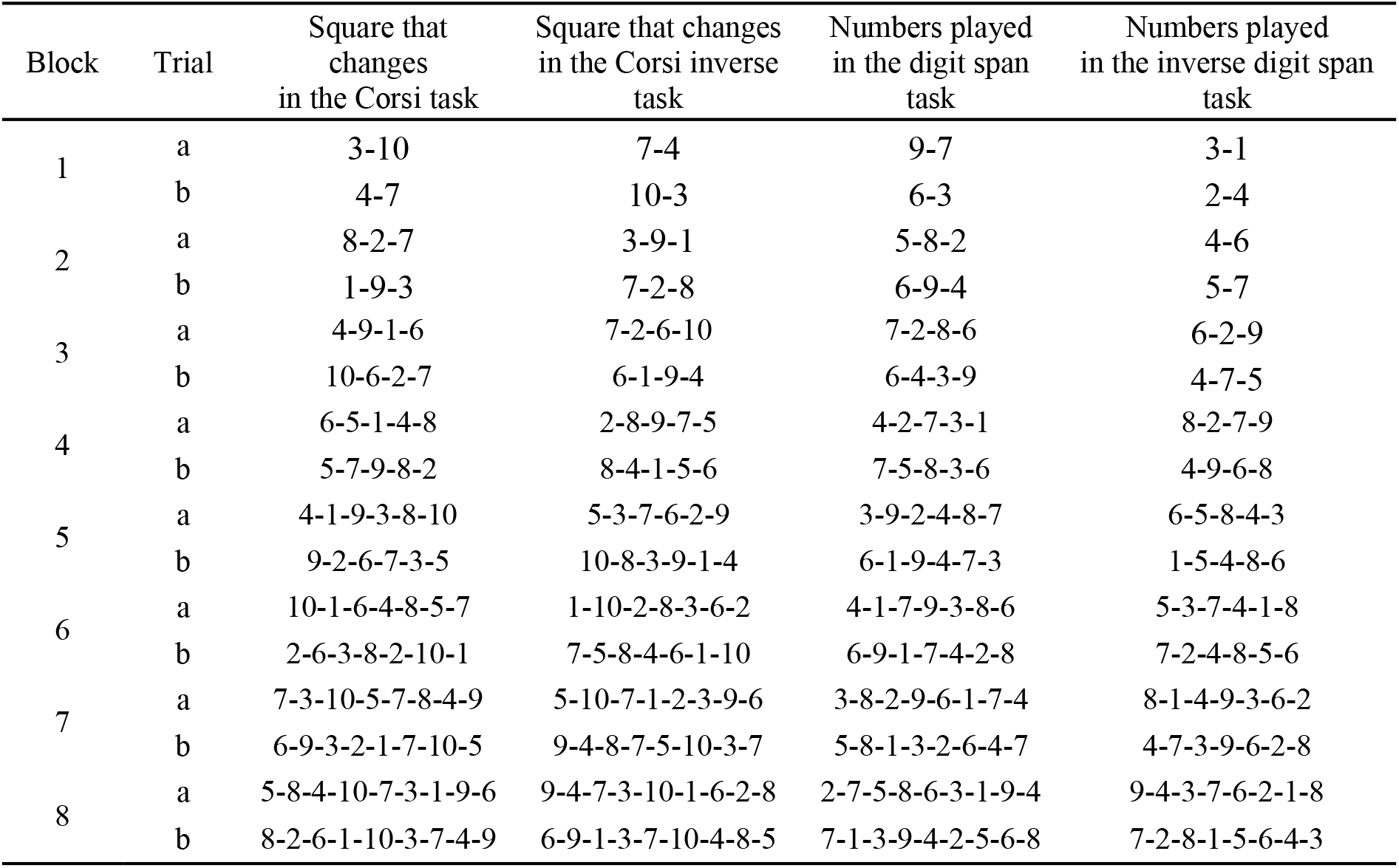
Stimuli of working memory tasks. Number of the square that changed in each of the sequences displayed in each trial of the Corsi and Corsi inverse tasks and the numbers played in each trial of the digits span and digit span inverse tasks. For the graphical display of the position of each square, see Fig 1.

For the <u>digit span</u> and the <u>inverse digit span</u> tasks, sequences of digits were presented to the participants. See Table 6 for a description of said sequences and the digits used in them. There were 8 blocks in total in the experiment, each of them containing two trials. In each trial, the participants were presented with a fixation point in the center of the screen while sequences of numbers were presented auditorily. Participants had to listen to the series of numbers that were presented at an approximate rate of one digit per second, and once each sequence finished, they were asked to repeat the numbers of the sequence out loud. For the <u>digit span</u>, this repetition had to be done in the same order as they listened to the numbers. For the <u>digit inverse span</u> task, they had to repeat them in the inverse order. The difficulty increased with each block, as the sequence length increased with each of them: the sequences of the first block consisted of two numbers, the sequences of the second block consisted of three numbers, and so on until the 8^th^ block, in which the sequences consisted of 9 numbers (see Table 6). After the two trials of each block, the experimenter indicated the amount of correctly retrieved sequences in that block. If the participant failed both of the trials, the experiment finished. If the participant repeated accurately one or two items, the experiment continued (see Fig 3 for a schematic representation of the experiment and see Table 6 for details on the trials).

**Fig 3:**
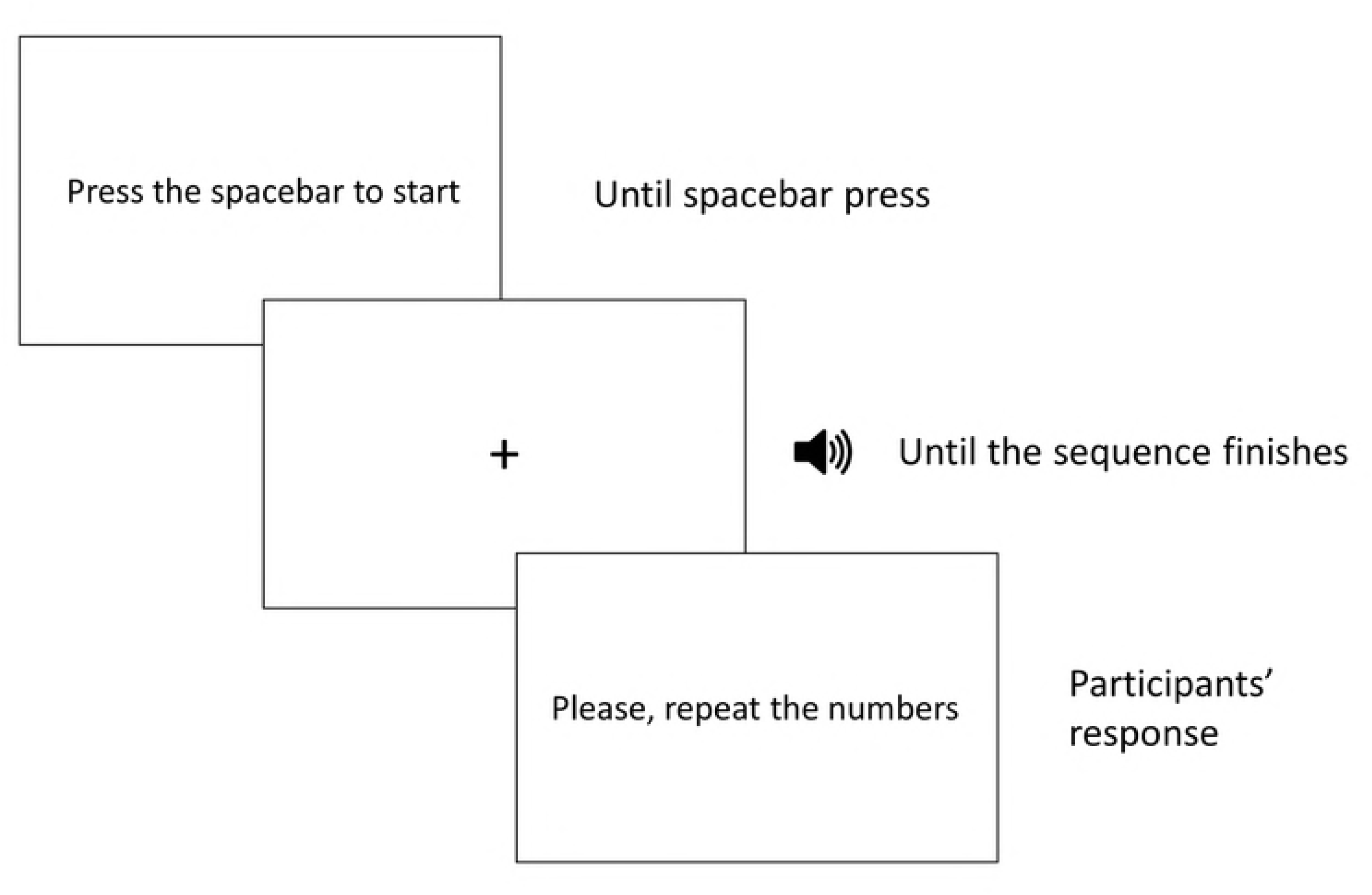
Schematic representation of the digits span and digits span inverse tasks.

#### Results

For the analysis, the amount of trials remembered out of 16 (2 trials in each of the 8 blocks) were compared between groups in each of the tasks. (see Table 7 for descriptive results). In the <u>Corsi task</u> Bilinguals remembered an average of 9.92 trials (SD=1.97) and monolinguals an average of 9.84 (SD=2.33), but the difference between the two groups was non-significant [*t*(178)=0.24, *p*>.81], and null hypothesis was also supported by the Bayesian Factor analysis (*BF*_01_=6.02). In the <u>inverse Corsi task</u> bilinguals remembered an average of 8.91 items (SD=1.67) and monolinguals an average of 7.98 (SD=1.95), and the difference between the two groups was significant [*t*(178)=3.45, *p*<0.01]. The alternative hypothesis was also supported by the Bayesian Factor analysis (*BF*_01_=0.03). In the <u>digit span</u>, bilinguals remembered an average of 8.89 trials (SD=2.16) and monolinguals an average of 8.6 (SD=1.92), but the difference between the two groups was non-significant [*t*(178)=0.95, *p*>.35], and the null hypothesis was also supported by the Bayesian Factor analysis (*BF*_01_=4.01). Finally, in the <u>inverse digit span</u>, bilinguals remembered an average of 8.97 items (SD=1.84) and monolinguals an average of 7.94 (SD=1.84), and the differences between the two groups was significant [*t*(178)=3.72, *p*<0.01]. The Bayesian Factor analysis supported the alternative hypothesis (*BF*_01_=0.01). The results in the Corsi and digit span asks were also analyzed using the highest recalled set size. Differences are not significant in the Corsi task (p>.43, Bilinguals recalled an average set size of 6.48, and monolinguals of 6.61). For the Corsi inverse, the recalled set size was significantly different for bilinguals (6.07 items on average) and monolinguals (5.67 items on average; p<.02). Following the same pattern, the recalled set size was not significantly different between groups in the digit span task (p>.59, bilinguals recalled an average size of 6.91 and monolinguals 6.82), but it was different in the backwards span task (p<.01), with bilinguals recalling slightly larger sets (6.92) than monolinguals (6.51).

**Table 7:** Results of working memory tasks. Mean number of items remembered and standard deviations (between parentheses) are displayed for each of the task of the working memory set of experiments.

|  | Monolinguals |  | Bilinguals |  |
| --- | --- | --- | --- | --- |
| Corsi | 9.84 | (2.33) | 9.92 | (1.97) |
| Corsi inverse | 7.98 | (1.95) | 8.91 | (1.67) |
| Digits span | 8.60 | (1.92) | 8.88 | (2.16) |
| Digits span inverse | 7.95 | (1.84) | 8.97 | (1.84) |

### Interim conclusion

The results obtained with the Corsi and the digit span tasks are complimentary and both show the same picture. In the standard forward version of the tasks, monolinguals and bilinguals reached the same level in terms amount of trials remembered. However, when their WM skills were pushed to the limit with the backward version of the task, bilinguals systematically outperformed monolinguals by remembering, on average, one item more, meaning that bilinguals were able to store, reverse, and produce sequences one step harder than monolinguals. While Bayesian tests favored the null hypothesis in both forward conditions, it unambiguously favored the alternative hypothesis when the backward tasks are analyzed.

### Additional analyses

To leverage our sample size and the extensive sociodemographic data in order to better understand how often unmatched socio-demographic factors or WM skills co-occur with significant bilingual advantage in EF tasks (i.e., the first four tasks), we conducted a systematic bootstrapping study. Firstly, we randomly sampled subsets of 25, 50, and 75 participants, 1000 times for each sample size, and measured how often the EF were significantly different between groups. Then, we explored how often the sociodemographic variables differed significantly in those samples where the EF differences were found.

With 1000 random samples of 25 participants, we only found a significant difference between groups in the flanker task in 2.1% of the samples, in 1.5% of the samples in the Simon task, in 7.3% of the samples in the numerical Stroop task and in 3.1% of the samples in the Stroop task. Out of those samples that showed a significant difference between groups, 9.46% showed a significant difference between group in Age (all of them produced by significantly older bilinguals), 4.05% of them showed a significant difference in IQ (83.33% of them produced by bilinguals having significantly higher IQ punctuations), and 25.68% showed significant differences in SES as measured by monthly incomes divided by household members (all of the significant cases were due to bilinguals scoring higher). Crucially, in 40.54% and 51.5% of these subsamples we also found a higher punctuation for bilinguals in the inverse Corsi and inverse digit memory tasks respectively. Regarding the EF tasks, 12.16% of the subsamples showed significant differences in the Flanker task (42.86% showing a bilingual advantage, and 57.14% showing a bilingual disadvantage), 12.16% of them showed significant differences in the Simon task (73.33% of them showing a bilingual advantage, 26.67% showing a bilingual disadvantage), 60.81% showed significant differences in the numerical Stroop (all of them indicating a bilingual advantage) and 22.97% showed significant differences in Stroop (83.87% indicating a monolingual advantage, 16.13% indicating a bilingual advantage).

With 1000 random samples of 50 participants, there were significant between group differences in the flanker task in 0.2% of the samples, in the Simon task in 0.3% of the samples, in the numerical Stroop task in 6.5% of the samples and in the Stroop task in 0.5% of the samples. Exploring only the samples that showed significant differences, 10.81% of them showed significant Age differences (all of them produced by significantly older bilinguals), 6.76% of them showed a significant difference in IQ (all of them due to bilinguals having significantly higher IQ scores), and 51.35% showed significant differences in SES (all of the significant cases were due to bilinguals scoring higher). Importantly, in 89.19% and 87.84% of these subsamples bilinguals outperformed monolinguals in the inverse Corsi and inverse digit span tasks. When it comes to the EF tasks, 2.70% of these subsamples showed significant differences in the Flanker task (all of them indicating a monolingual advantage), 4.05% of them showed significant differences in the Simon task (66.67% of them showing a bilingual advantage, 33.33% showing a bilingual disadvantage), 87.84% showed significant differences in the numerical Stroop (all of them indicating a bilingual advantage) and 6.76% showed significant differences in Stroop (all of them indicating a monolingual advantage).

With random samples set to 75 participants, and thus closer to our actual number of participants, there was no sample comparison that showed significant differences between groups in the flanker, the Simon or the Stroop tasks. In 2.6% of the cases a significant difference in the Numerical Stroop task was found, where bilinguals performed better than monolinguals. In those cases, 3.85% of the cases showed significant IQ differences (all of them produced by significantly higher punctuations obtained by the bilingual sample), and 88.46% showed significant differences in SES (all of the significant cases were due to bilinguals scoring higher), and the 100% showed a better performance of bilinguals in the inverse Corsi and inverse digit span tasks.

Additionally, systematic multiple regression analyses were conducted to try to capture any possible influence of the sociodemographic factors in the indices obtained in each of the tasks. First, three models were built for each of the four EF tasks task. The Stroop or conflict effect (in milliseconds) was used as the dependent variable, and Age, IQ punctuation (correct responses in the abridged version of the K-Bit task used in the experiment), SES (the value resulting from dividing monthly income by household members) and LexTale punctuation in Spanish were included as independent variables in Model 1. Model 2 also included Group (Bilinguals and Monolinguals) and Model 3 included the interaction terms between Group and the rest of the demographic and linguistic factors. None of the models explained enough of the variability in the data (all R^2^ < .06) and none of them reached significance (all *p*s> .17). Importantly, in none of those models was Group a significant predictor (all *p*s> .3) nor was a significant interaction with the rest of the sociodemographic factors (all *p*s>.17).

The same regression approach was used to explore the possible influence of the sociodemographic factors in the WM punctuations. The number of correctly recalled items was used as the dependent variable, and Age, IQ, SES and LexTale scores in Spanish were included as independent variables in Model 1. Model 2 also included Group (Bilinguals and Monolinguals) and Model 3 included the interaction terms between Group and the rest of the demographic and linguistic factors. For the <u>Corsi task</u>, Model 1 resulted in a significant regression equation [*F*(4,175)= 2.98, *p*<.03], with a R^2^ of .06. Model 2 was significant as well [*F*(5,174)= 2.40, *p*<.04] with a R^2^ of .06, but did not significantly improve the first model (*p*>.74). Model 3 was not significant (*p*>12) and did not improve the model either (*p*>.70). Thus, following Model 1, participants’ predicted Corsi punctuation is equal to 11.75 - 0.10 (age) + 0.16 (IQ). While IQ was a significant predictor (*p*<.01) and Age was marginally significant (*p*<.06), SES and Lextale scores were not (all *p*s>.30). Group and interaction between Group and other factors were not significant predictors in any of the models (all *p*s>.29). For the <u>Corsi inverse</u> task, Model 1 was a significant regression equation [*F*(4,175)= 3.01, *p*<.03], with a R^2^ of .06. Model 2 was significant as well [*F*(5,174)= 4.51, *p*<.01] with a R^2^ of .12, and significantly improved the first model (*p*<.01). Model 3 was also significant (*p*<.01) but did not improve the model (*p*>.74). Thus, following Model 2, participants’ predicted Corsi inverse punctuation is equal to 2.22 + 0.09 (IQ) +0.06 (LexTale) + .85 (Group). Participants’ recalled inverse digit items increased 0.09 for every increase in punctuation in the IQ test, 0.06 for every increase in punctuation in the Spanish LexTale task, and bilinguals recalled .85 items more than monolinguals. Punctuation in LexTale was a significant predictor (*p*<.05), as well as Group (*p*<.01), and IQ was marginally significant (*p*<.08). SES and Age were not (all *p*s>.70). The <u>digit span</u> task followed a similar pattern as the one showed by the Corsi task: Model 1 was a significant regression equation [*F*(4,175)= 3.18, *p*<.02], with a R^2^ of .07. The inclusion of Group (Model 2) produced a significant model [*F*(5,174)= 2.73, *p*<.04] with a R^2^ of .07, but did not significantly improve the first model (*p*>.33). Model 3 was significant as well [*F*(5,174)= 2.07, *p*<.04] but did not improve the model either (*p*>.30). Thus, following Model 1, participants’ predicted Digit Span punctuation is equal to 6.18 - 0.12 (age) + 0.12 (IQ). IQ and Age were significant predictors (*p*s<.03), SES and Lextale were not (all *p*s>.16). Group and interaction between Group and other factors were not significant predictors in any of the models (all *p*s>.12). Resembling what was found for the Corsi inverse task, the analysis for the <u>inverse digit span</u> task showed that Model 1 resulted in a significant regression equation [*F*(4,175)= 5.07, *p*<.01], with a R^2^ of .10. Model 2 was significant as well [*F*(5,174)= 6.90, *p*<.01] with a R^2^ of .17, and significantly improved the first model (*p*<.01). Model 3 was also significant (*p*<.01) but did not improve the model (*p*>.54). Thus, according to Model 2, participants’ predicted Corsi inverse punctuation is equal to 2.38 - .10 (Age) + 0.14 (IQ) +0.05 (LexTale) + 0.96 (Group). Participants’ recalled inverse digit items decreased for .1 for every older year, increased 0.14 for every increase in punctuation in the IQ test, 0.05 for every increase in punctuation in the Spanish LexTale task, and bilinguals recalled .96 items more than monolinguals. As it can be seen, the significant predictors were Age (*p*<.03), IQ (*p*<.01), and Group (*p*<.01), and Lextale was marginally significant (*p*<.07). SES was not (all *p*s>.17). In this second set of tasks, i.e. the WM tasks, we did not conduct a detailed bootstrapping analysis, because during it, we observed that the majority of the subsamples showed a bilingual advantage in the inverse versions of the WM tasks (>42% of the 25 sample subsets, >83% in the 50 sample subsets, and >99% in the 75 sample subset), and thus the co-occurrence with other factors would not be very informative.

### Interim conclusion

Bootstrapping analyses indicated that the bilingual advantage in random samples was much more frequent in small rather than in big sample sizes, and also that the advantages in EF tasks co-occurred mostly with significant differences in SES and WM tasks favoring bilinguals, followed by Age and IQ differences. Unfortunately, and probably due to the low variability of said factors due to our matching procedures, the models used to further explore this issue with multiple regression analysis did not reach significance. On the other hand, multiple regression analysis conducted for WM tasks indicated that some sociodemographic factors, mostly Age and IQ, predicted participants’ performance in the forward WM tasks irrespectively of their linguistic profile. On the contrary, reverse WM task punctuation was also predicted by the linguistic profile and by the Spanish proficiency.

### General Discussion

This study aimed at exploring the potential effects of bilingualism on two main processes: EF and WM. To the best of our knowledge, this is the first study in which large samples of bilinguals and monolinguals are extensively tested using multiple tasks to asses both their EF abilities (tasks 1 to 4) and WM span (tasks 5 to 8) while relevant demographic factors (age, IQ, SES, educational level and immigrant status) are controlled for.

The first hypothesis put to test was the enhancement of EF as a consequence of bilingualism. In tasks 1 to 4 we attempted at verifying the reliability of the bilingual advantage hypothesis by using the same tasks and equivalent populations that the previous studies did (13,19,44,45) but attempting to account for the concerns raised by the criticisms to these studies (26,30,31). In that regard, the predictions were rather straightforward. If bilingualism provides an advantage in EF independently of the effects of the controlled factors, bilinguals would have shown a reduced *conflict* or *Stroop* effects when compared to monolinguals (8) or faster global reaction times (25). The flanker, Simon, Stroop and numerical Stroop tasks produced the expected classic patterns, with strong and constant conflict effects in all of them, mainly driven by the incongruity effect. Each condition (incongruent, congruent and neutral) behaved as expected and in accordance with preceding literature. However, none of the effects or conditions varied significantly across language groups, and language groups did not overall differ in reaction time either. Furthermore, the Bayesian factor analysis clearly showed that the null hypothesis was the most suited explanation for the results we obtained: bilinguals and monolinguals did not differ as to how they face the demands of changing tasks with congruent and incongruent trials. Importantly, the results obtained using bootstrapping analyses shed additional light on the role of the uncontrolled socio-demographic factors. In the subsamples where a significant EF differences between groups were found, they very often co-occurred with some unmatched sociodemographic factor, especially SES. The impact of SES, both together with and independently of bilingualism, has been gaining researchers’ attention lately. For example, Hartanto, Toh, and Yang (85), reported that both high levels of SES and bilingualism correlated with better EF in children, but only SES was a reliable predictor of verbal WM. Importantly, they found that bilingualism predicted advantages in EF tasks in low SES groups only. Altogether, these results provide credibility to the concerns that the bilingual advantage obtained previously in these tasks might be found as a consequence of unmatched external factors, rather than by bilingualism itself (30), and that it disappears when the confounding factors are controlled for (26,31,46–49). Along the same lines, see a very recent meta-analysis of the effect sizes found in 152 different studies, where no strong support of the bilingual advantage in conflict monitoring, inhibiton or WM is found (53).

Far from ending, though, the debate around the bilingual advantage feeds from growing evidence pointing towards both directions. Our data here, together with the recent findings of no bilingual advantage in Basque-Spanish bilingual children (46,47) and seniors (49) addresses whether this advantage in EF appears in truly bilingual speakers in a truly bilingual community. And the answer this far, although it can only be extrapolated to comparable populations and societies, is a robust no. The conclusions drawn from the analyses conducted in the present article can only be circumscribed to a very specific kind of bilingual population, yet as important as any other – balanced and native bilinguals immersed in a bilingual society, but we believe that they are of crucial importance to better understand and reframe the current perspectives on the bilingual advantage debate. In the Basque society, the ratio of usage of Basque and Spanish differs between age groups, regions and social spheres, being on average 20% of the citizens older than 16 years who use Basque as much as or more than Spanish (according to Basque Institute of Statistics, *Eustat*), and thus creating an heterogeneous linguistic mosaic in which language use varies widely across social groups.

Therefore, we wonder whether the use of the EF that bilingualism asks for is strong enough to provoke changes at the behavioral level. The argument for a bilingual advantage on executive control tasks rests on the idea that monolinguals do not switch between two languages, since they only have one available. However, all human beings face situations in which they have to inhibit salient responses constantly and monitor the environment, in both general social situations and when performing concrete actions. For example, people do switch between comprehension and production when they talk to somebody, they do switch and keep their monitoring abilities strongly activated when they have to drive and talk to somebody, or they inhibit salient responses when they have to adapt their speech and manners to different social situations, which can range from casual to very formal. Thus, monolinguals also efficiently use their switching, inhibitory and monitoring skills, and it is unclear whether language switching in bilinguals imposes a heavier burden than the one imposed to everyone, monolingual or bilingual, in their daily life. This interpretation follows the assumption made by the bilingual advantage hypothesis, namely, that language control and general executive control functions are two completely overlapping mechanisms (e.g. “Crucially, the mechanism that reduces attention to the non-relevant language system is the same as that used to manage attention in all cognitive tasks”, 14, p.41), implying that EF are domain-general and that they apply to every situation in which they are needed, linguistic or not. Hence, training in one concrete aspect directly implies an improvement in any other context where the same EF are needed. However, a different interpretation comes from questioning the roots of the advantage: what if the EF were not as domain general as they have been claimed to be? If they were, the performance in the four EF tasks used in this study should correlate with each other, inasmuch as they are supposed to reflect the same general ability. Our results clearly show that they do not, indicating different underlying mechanisms (in this same regard, see 30,64). The degree of domain-specificity of the different EF components - especially switching and inhibition - has been recently questioned and tested using both behavioral and neuroimaging measures, and results tend to indicate that the domain-generality assumption is debatable at best (86–89). Behaviourally, the performances in linguistic and non-linguistic switching tasks do not correlate with each other (e.g.,86,87,89), similarly to linguistic and non-linguistic inhibition tasks (such as the n-2 task, see (90). While it seems that there is a strong overlap in the brain areas responsible for linguistic and non-linguistic switching, whether or not domain-general inhibition and language control respond to the same brain mechanisms is still unclear(86,88,91,92). Furthermore, and even in the very same field of language control, the most recent studies speak of different language control mechanisms relying on different neural substrates when applied to language comprehension and production (93). If the domain-specificity of the EF is true, it would invalidate the training transfer assumption that the bilingual advantage hypothesis is based on.

On the other hand, the second hypothesis tested in the present article predicted a potential bilingual advantage in WM skills. We observed that bilinguals outperformed monolinguals in the backward versions of both the Corsi and the digit span tasks, with no differences in the forward versions. As opposed to the EF, this cognitive ability might have a stronger domain-general component: even though some authors have argued for separate WM stores and mechanisms for different sensory domains (domain-specific WM, 94,95), others defend that the maintenance system that retains the stimuli is unitary (the domain-general perspective, (61,96,97) despite the existence of domain-specific stores of the WM. Using neuroimaging techniques, some authors found different brain regions involved when processing the stimuli from different domains (98–100) while some others found the same region involved in memory maintenance no matter the domain (101–104). Trying to solve this issue, Li et al. (105) found functional networks responsible for domain-general and domain-specific processes in WM. Interestingly, while specific networks showed an important role only during encoding, domain-general networks showed load-dependent patterns during encoding, maintenance and retrieval. Importantly, in our data (tasks 5-8), the differences were found only in tasks that involved a more complex processing and retrieval (transforming the encoded information to the backwards series) of the information stored, i.e. in the backward conditions (see also 74; for a bilingual advantage in more demanding memory tasks but not in simple ones; and 106; for results showing a bilingual advantage in inverse digit span tasks). Precisely, the situations in which the domain general WM system would be required (105). Unlike the domain-specific networks, not susceptible to training transfer, domain-general WM abilities are capable of improvement via enhancement of some different domain (like bilingualism), and the existence of a transfer is worth considering as it has been shown that training can improve WM (see 55–58, but see also 59, for evidence against the beneficial effects of training in WM).

An interesting twist to this hypothesis comes from the fact that both EF and WM are strongly related (107), and differences in EF tend to correlate with differences in WM (61). This is especially prominent in demanding WM tasks that require storing and processing of information (62). More specifically, and as it was mentioned in the introduction, the relation between WM and updating ranges from related but separable (64) to equated (23). In turn, it has been argued in the bilingual advantage literature that said advantage stems from improved monitoring (i.e., updating) abilities, which are captured in the classic EF tasks as faster overall reaction times (25). This triple equation needs to be disentangled to further understand the source of the differences found in the present paper and in the bilingual advantage literature. Some authors have argued that it is in the WM where the bilingual advantage is located, and then, due to its close relation to monitoring, this advantage reflects in EF tasks (65). Arguably, considering the close relationship between the two constructs, this could be counter-argued by attributing the source of the advantage to EF abilities which then translates to an indirect improvement of WM. For example, Morales, Calvo and Bialystok (74) report a bilingual advantage in WM tasks only when the EF demands imposed by the task are high, and therefore they argue that it is the role of EF that improved bilinguals’ performance in WM (along the same lines, see 106). However, note again that the validity of these results is put to question by the lack of control of several factors, such as ethnicity (the group of bilinguals is formed of individuals with more than 15 different second languages, indicating significant linguistic and probably ethnical differences) or SES (just parents’ educational level is reported, and very scarcely. To our understanding, the results in the present study picture the opposite: the performance in the EF tasks was similar for bilinguals and monolinguals, indicating no bilingual advantage for our sample of native balanced bilinguals immersed in a bilingual society. On the other hand, we consistently found a bilingual advantage in the backward WM tasks where information has to be actively processed and retrieved in a complex way. In principle, the absence of an advantage in EF could be arguably due to the ceiling effect that adults of this age feature in EF abilities (108) that prevents any potential enhancement in EF from being captured. However, when random resampling analyses were conducted for a thousand times for each different sample sizes, the bilingual advantage was found in some variable percentages of the cases, so there was still room for differences. Interestingly, the sets that showed a bilingual advantage also displayed an advantage in backward memory tasks in the majority of the cases, as well as other unmatched factors such as SES. This dissonance makes us hypothesize that bilingualism does improve WM abilities, and then this can –but does not necessarily have to – translate into an enhancement of EF abilities when interacting with other factors, but not the other way around. Had the advantage in memory been a consequence of EF functions, the tasks employed to measure said functions (i.e., tasks 1-4) should have shown an advantage in some of the dimensions as well. Furthermore, our findings show no general advantage in monitoring –i.e., faster RTs– but better WM abilities. This supports the idea that these two constructs, even though often equated (23), might overlap but are different (64). It also strengthens the argument made by Namazi & Thordardottir (65) after they found that a WM advantage in bilinguals might potentially lead to an apparent advantage in EF tasks as well, and thus WM should be held constant to explore EF abilities.

All in all, the results obtained from the 8 tasks conducted in the present study show a very stable pattern. Firstly, native and balanced bilingualism does not improve bilinguals’ general EF abilities when compared to carefully matched monolingual counterparts. Secondly, bilingualism improves WM when the task requires an active and complex processing and retrieval of the encoded information. This pattern is interpreted as a consequence of the domain-specificity of the EF and the encoding processes of WM, and is thus not susceptible to be indirectly trained. The load-dependence of the maintenance and retrieval of the encoded information in WM tasks has been shown to be domain general. This makes the backward conditions of the memory tasks suitable for improvement due to training transfer. Despite the information provided by the lack of correlation between the EF tasks, we did not collect any data testing other aspects of EF abilities, and therefore this interpretation of the results is rather speculative.

As a general consideration, it should be born in mind that the conclusions derived from this study are generalizable only to the populations and situations similar to the ones tested here, that is, lifelong, native and balanced bilinguals (in particular the case of Basque-Spanish bilinguals, see (46,47,49). When different bilingual profiles are considered, the same patterns are not completely guaranteed. For example, the factors of immigration and late bilingualism should be specially considered. Immigration usually involves moving to a different language-speaking country and it forces people to become bilingual, so it often co-occurs with late bilingualism and both factors can be confounded when the significant effects of bilingualism are explored (see 12,13,28,44,109,110, among others, for studies reporting bilingual advantages that tested bilingual samples formed by mostly immigrant individuals). Immigrants, who generally happen to be bilinguals, would show some enhancements maybe wrongly associated to bilingualism when compared to non-immigrants, who happen to be monolinguals. It is still unclear whether those effects are purely produced by a late bilingualism, by being an immigrant, or a combination of both. It seems coherent to propose that native bilingualism does not necessarily bring any eventual benefit, simply because native bilingualism does not imply a strong cognitive effort to deal with two languages from birth, and there is no strong reconfiguration needed to incorporate them into the mental repertoire. Training and cognitively demanding acquired skills that lead to an enhancement of attentional skills (111,112) are found prominently when the mentioned training happens late in life (113). Similarly, lately acquired bilingualism would indeed require the new bilinguals to re-adjust their mental repertoires to be able to accommodate the existing system to the newly acquired language. This cognitive effort in adapting the system could lead to an improvement in the EF. The way in which a late acquisition of a second language would change the ongoing developmental trend of different high level cognitive skills (among others, the EF) is a venue worth exploring for future research looking for any kind of bilingual advantage (see 114, for authors arguing for a stronger bilingual advantage when the second language is acquired early; see but see 115,116, for evidences favoring an advantage when L2 is acquired late in life, and see 18 for no differences between monolinguals, early bilinguals and late bilinguals).

### Conclusions

The results from this set of tasks suggest that bilingualism is not enough to enhance young bilingual adults’ EF skills relative to the ones of young monolingual adults. Both groups behaved similarly in all the tasks, as measured by the different indices and conditions which, importantly, did not correlate across tasks. Thus, it is argued that EFs are not as domain general as they were believed to be, and therefore the hypothesis of a training transfer produced by bilingualism is not supported. The results of the bootstrapping analysis indicate that when the bilingual advantage in EF is found, it very often co-occurs with significant differences in socio-demographic factors and memory abilities, suggesting that previous findings might have been a consequence of unmatched factors.

From the results from the WM tasks we clearly see that there was no effect of bilingualism in the easiest versions of the tasks (i.e., forward versions where only storing and repeating is needed) but it does improve the WM skills required in backward tasks where storing, manipulation and retrieval are used. We interpreted this selective bilingual advantage as based also in the domain-specificity of some abilities. Previous findings have shown that encoding relays on domain-specific WM, and therefore no training transfer would be expected. However, the backward task has a stronger component of maintenance of information, manipulation, and retrieval, which have been shown to be more domain-general, and consequently more susceptible to training transfer.

The practical contributions of this work are twofold. Firstly, it is the first time that a bilingual advantage is found in WM tasks in carefully matched large sample sizes of balanced and native young adult bilinguals immersed in a bilingual society. Secondly, it emphasizes the need of the methodical sample matching, since we found no bilingual advantage in EF when samples were matched for known confounding factors. Importantly, the majority of the subsamples that showed a significant bilingual advantage in the bootstrapping analysis co-occurred with differences in other sociodemographic factors.

Altogether, the different analysis conducted with the current data and the previous findings of the field lead us to conclude that the previous findings of a bilingual advantage in EF might have been a product of the uncontrolled non-linguistic characteristics of the cohorts of participants tested. Instead, bilinguals seem to benefit from their idiosyncratic language context in situations where an active use of the elements in the WM is needed, and it is in this context when they outperform their monolingual peers. We believe that the results shown here will help to reinterpret the theories behind the bilingual advantage theory and to narrow down the scope in future research to help identifying the critical factors that make the bilingual advantage to show up sometimes.

## Acknowledgments

For this research, JAD has been partially funded by grant PSI2015-65689-P from the Spanish Government (http://www.ciencia.gob.es/), EA has been partially funded by grant PI2015-1-27 from the Basque Government (http://www.euskadi.eus/gobierno-vasco/departamento-educacion/), and MC has been partialy funded by ERC-AdG-295362 grant from the European Research Council (https://erc.europa.eu/), by the AThEME project funded by the European Union (grant number 613465, http://www.atheme.eu/), and by grant Centro de Excelencia Severo Ochoa SEV-2015-0490 provided by the Spanish Government (http://www.ciencia.gob.es). The funders had no role in study design, data collection and analysis, decision to publish, or preparation of the manuscript..

